# Association of intrinsic functional connectivity between the locus coeruleus and salience network with attentional ability

**DOI:** 10.1101/2022.11.01.514703

**Authors:** Joshua Neal, Inuk Song, Benjamin Katz, Tae-Ho Lee

## Abstract

The LC is a brainstem region associated with broad physiological and neural arousal as part of the release of norepinephrine, but it has increasingly been associated with multiple specific cognitive processes. These include sustained attention, deficits in which are associated with a variety of neuropsychological disorders. Neural models of attention deficits to date have focused on interrupted dynamics between the salience network (SAL) with the fronto-parietal network (FPN), which has been associated with task-switching and processing of external stimuli, respectively. Conflicting findings based on these regions suggest the possibility of upstream signaling leading to attention dysfunction, and recent research suggest the LC may play this role. In this study, resting-state functional connectivity (FC) and behavioral performance on an attention task was examined within 584 individuals. Analysis revealed significant clusters connected to the LC activity in the bilateral insula, anterior cingulate cortex (ACC), and bilateral ventral striatum, all regions associated with the SAL. Given previous findings that attention deficits may be caused by dysfunctions in network switching by the SAL, our findings here further suggest that dysfunction in LC signaling to the SAL may interfere with attention.

## INTRODUCTION

The locus coeruleus (LC), a small region in the brainstem, is associated with multiple aspects of attention. The LC’s association with attentional processes is most often cited as part of its’ role in the release of norepinephrine, a neurotransmitter known to cause broad general arousal within the brain (Nai-shin & E, 1973). Beyond broad release for general arousal, compelling evidence suggests that the LC may produce more specific signaling, such as those related to the activation or deactivation of specific networks (Devilbiss & Waterhouse, 2011; Jacobs et al., 2015) or tasks (Clewett et al., 2018; Lee et al., 2018; Vazey et al., 2018). LC signaling-related modulation could then be relevant to the collection of functional connectivity networks associated with higher order cognitive processes, including attentional control.

Models for understanding neural activity in the human brain establish common networks through functional connectivity (FC), defined as the interrelationship between neuronal activity in different brain regions (Aertsen et al., 1989; Friston et al., 1993) and most commonly calculated as the correlation of regional activation time series when observed in a resting state (Lowe et al., 2000). The neural substrates of dysfunction in sustained attention can, like those of other cognitive processes, be understood through the triple network model (Menon, 2011). Within this model abnormal signaling relationships between the dorsolateral prefrontal cortex (DLPFC), a core region of the fronto-parietal network (FPN), the salience network (SAL), including the anterior insula and anterior cingulate cortex (ACC), and the default mode network (DMN), including medial prefrontal cortex (mPFC) are understood as neural origins for cognitive dysfunctions across pathologies (Wang et al., 2020; Wei et al., 2016; Yu et al., 2019).

Given the support for larger, systems-level brain networks (Buckner & Vincent, 2007), as well as interactions between them, it is necessary to consider the possibility of extended regions and networks that may be related to attention deficits. The SAL would be a suitable candidate for further analysis, having been 1) identified within the triple network model as a mediating network between the LC and other networks and 2) demonstrated to direct differential neural activation to either the FPN or DMN (Goulden et al., 2014). Inattention may reflect dysfunction within multiple attention constructs, but sustained attention (the capacity to maintain a primary focus while simultaneously processing external stimuli for unusual/undesired effects) relates most directly to common functional needs associated with attention (Sarter et al., 2001). A neural explanation of sustained attention dysfunction (see Fortenbaugh et al., 2017) would suggest abnormalities in network switching or the mediation of differential neural activation to appropriate networks relevant to given stimuli, inherently implicating the SAL within this model.

While shifting the focus of analysis from the FPN and DMN to a switching network upstream, such as the SAL, may prove valuable to discern the aforementioned FC findings, it’s worth inquiring further into the possible causal mechanisms for the SAL to do this. This study is based on one such possibility: that LC communication with the SAL may itself be the causal mechanism for SAL network switching. On the neurotransmitter level, the β-adrenoreceptor blockade of LC activity also coincides with a decrease in SAL activation (Dahlöf et al., 1981). Within animal models, electrical and optogenetic stimulation of the LC changes its tonic and phasic activity, which in turn rapidly reconfigures the neural dynamics between various intrinsic networks including the SAL(Devoto et al., 2005; Jodoj et al., 1998; Marzo et al., 2014; Zerbi et al., 2019). Similarly, a recent conceptual account of attentional processing suggests that the locus coeruleus – noradrenergic (LC-NA) system, along with the SAL, may play a key role in allocating attentional resources for processing selectivity and maintenance to guide appropriate behavioral responses (Sara, 2009).

Thus, research from other modalities and theoretical perspectives support the possibility that the LC-SAL connection may play an important role in instigating attentional processing and sustaining selective attention within the brain. Explaining this in terms of attention processing dysfunction, the SAL failing to interact properly with the LC and thus not favoring connectivity to externalizing networks, such as the FPN, may lead the SAL to fail at switching primary connectivity status away from internalizing networks, primarily the DMN. One possible scenario is that the SAL is more likely to be influenced by the LC in establishing connection primacy with other intrinsic networks, and therefore setting processing priorities in executive functions afterward. For example, a recent study demonstrated that older adults, who are generally more prone to distractors, showed decreased LC-SAL connectivity compared to younger adults, but there was no direct connectivity between the LC and FPN (Lee et al., 2018). However, an alternative explanatory mechanism could be that the SAL, receiving feedback from the cortex and its intrinsic networks, signals to the LC to alter arousal function within those networks, which the LC could then do as part of the LC-NA system. Unsworth and Robison’s review (2017) identifies top-down attentional processing as being related to working memory performance, and specifically noted, as part of their model, the ability of the FPN to signal back through the SAL. This is identified as being related chiefly to the SAL’s role in directing neural signaling to other networks, with the subsequent FPN-DMN anti-correlation being associated with cognitive processes such as working memory (Keller et al., 2015), and damage to the SAL being related to impairments in DMN deactivation (Bonnelle et al., 2012). Direct neural patterns from the FPN and SAL to the LC have been previously identified (Jodoj et al., 1998; Rajkowski, 2000), and a model of LC-NA activation with an optimal degree of signaling (Aston-Jones & Cohen, 2005) would appear to support FPN down-regulation of the LC through the SAL in the absence of stimuli. The potential importance of the SAL responding to such negative feedback should be understood through the greater context of holistic network function. Just as excessive DMN activation may lead to inattention though a relative lack of resources for processing external stimuli via the FPN, excessive FPN activation in the absence of external stimuli may support inattention towards self-oriented tasks, such as memory processing. The SAL’s role in network switching and signaling is vital for the proper regulation of other intrinsic networks, so that each may function within a specific level of appropriate activation.

Furthermore, substantial evidence has been found connecting the various elements of this model to attentional processes and inattentive psychopathologies. The LC has been repeatedly linked to cognitive arousal, central executive tasks, and a wide variety of pathologies. Altered LC functional connectivity to sensorimotor networks has been linked to motor deficits in autism spectrum disorder (ASD) (Huang et al., 2021). The LC may also mediate hyperresponsivity symptoms in post-traumatic stress disorder (PTSD) patients (Naegeli et al., 2018). Additionally, LC neurodegeneration has consistently been identified as an initial biomarker within Alzheimer’s Disease, as well as other neurodegenerative conditions (Matchett et al., 2021). Even within a non-pathological context, the LC’s signaling discrepancies have been associated with inattentive symptoms in older adults (Lee et al., 2018).

Thus it is worthwhile to examine the association between LC-SAL connectivity and attention performance. To examine the intrinsic functional connectivity between the LC and SAL, an analysis utilizing resting-state fMRI was performed to best isolate the average functional connectivity from any stimuli or task-based activation (Biswal et al., 1995). LC-SAL functional connectivity was defined by the correlation of activation between the LC and the anterior insula, which is the region of interest (ROI) most consistently associated with the core of the SAL (Uddin, 2015). Our behavioral observation was the grand mean performance on the Attention Network Task (ANT), a neurocognitive test for alerting, orienting, and executive attention (Fan et al., 2002). We hypothesized that LC-SAL functional connectivity should be positively correlated with ANT attention performance.

## METHOD

### Participants

The present study utilized resting-state fMRI data from the enhanced Nathan Kline Institute (*NKI*)-Rockland project (Nooner et al., 2012; Tobe et al., 2022), which was initially downloaded via the Mind Research Network’s collaborative informatics and neuroimaging suite (*COINS*; Landis et al., 2016). Participants were initially selected for inclusion of all primary variables and covariates, including age, sex, the attention network test score (ANT; Fan et al., 2002), MRI data with full-coverage of both T1 and EPI, and to include only those without severe motion (mean framewise displacement, FD > 0.5mm). This resulted in the inclusion of 584 individuals (mean age = 39.5 years, SD = 20.6, SEM = 0.85, range = 8-83 years; 63.4% female). All fMRI data were collected with a 32-channel head-coil for T1 image (T1-MPRAGE; TR=1950 ms; TE = 2.52 ms; FA = 9°; 1-mm isotropic voxel; FOV = 256 mm) and echo-planar image (EPI; 364 volumes; 2-mm isotropic voxel; 64 slices; TR = 1400 ms; TE = 30 ms; FA = 65°; matrix size = 112 × 112; FOV = 224 mm).

### Attentional behavior measure

To measure an individual’s attention ability, we used the Attention Network Test score (ANT), a neurocognitive task designed to evaluate attentional performance (Fan et al., 2002). The task requires participants to correctly identify which direction a center arrow within a row of identical arrows is pointing, with the surrounding arrows either all pointed in the same direction (congruent) or in the opposite direction to the center arrow (noncongruent). Additionally, prior to each trial one of three cue conditions would be provided: no cue, a central cue, or a spatial cue. The combination of congruent and non-congruent conditions with these cues are designed to test individual attentional networks, and individual scores for alerting, orienting, and executive attention are calculated. However, the independence of these three scores has been repeatedly questioned (Galvao-Carmona et al., 2014; MacLeod et al., 2010; McConnell & Shore, 2011), especially within single measures of the task for participants. Use of a grand mean across conditions has been previously established as a means to assess a more general and integrated level of attentional processing (e.g., Johnson et al., 2008; van den Hurk et al., 2010). Thus, in our study, we adopted the grand mean performance score as the primary behavioral attention measure (mean = 617.43, SD = 119.91, SEM = 4.96, range = 410 – 1272).

### fMRI preprocessing

For the MRI preprocessing, we used FMRIB Software Library (FSL) with ICA-AROMA (Pruim et al., 2015) and Advanced Normalization Tools (ANTs; Avants et al., 2014). For both T1 and EPI, skull stripping and tissue mask segmentation (CSF/WM/GM) after bias-field correction for T1 images were conducted. The EPI images were then preprocessed including first 10 volumes cut, motion correction, slice-timing correction, intensity normalization, regressing out CSF/WM with individually segmented masks, ICA denoising using ICA-AROMA (Pruim et al., 2015), and registration to standard MNI 2-mm brain template.

### LC seed functional connectivity

Based on our hypothesis, we conducted a seed-based connectivity analysis using the LC as a seed region. We first transformed a standard structural LC mask (voxel number k = 160; Keren et al., 2009) to each individual’s native space. we then extracted the time-series of LC activity from the individual’s preprocessed EPI. Importantly, the LC seed activity was extracted from non-smoothed (Alakörkkö et al., 2017) and ICA-based physio corrected EPI images on the native space to increase LC signal fidelity (Lee et al., 2018; Prokopiou et al., 2022). Finally, group-level whole brain connectivity analysis was conducted to examine changes of LC connectivity with attention ability (i.e., ANT score) based on the individual LC seed connectivity map. We then used the FSL’s ‘*randomize*’ function (5000 permutations; Winkler et al., 2014) combined with threshold-free cluster enhancement correction (TFCE; Smith & Nichols, 2009) at corrected-*p* = .05 and 8-mm smoothing. In the whole-brain multiple regression model, we included gender, age and motion (mean FD) as nuisance regressors. Given the previous findings that LC connectivity changes quadratically with age (e.g., Song et al., 2021), the squared age was included in the model.

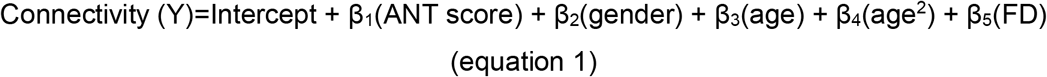

### Salience network, attentional executive region and attention performance

In addition to the LC-seed connectivity analysis that showed the LC-SAL connectivity is associated with the ANT score (see Result), we further examined the role of the FPN in LC-SAL dynamics, as studies have demonstrated that the dorsolateral prefrontal cortex (DLPFC), the core region of FPN, is involved in attention and/or executive functions (Jung et al., 2022; Laird et al., 2011; Luna et al., 2020; Nejati et al., 2018; Niendam et al., 2012; Thiele & Bellgrove, 2018) based on the interaction with the SAL (Chand & Dhamala, 2017; GÜrsel et al., 2018). To this end, we extracted time-series of the SAL and DLPFC regions. For the time-series of the SAL, we extracted the signal within the significant voxels of the LC-SAL connectivity results (see Figure 1). For the DLPFC, we used the automated anatomical labeling 2 atlas (AAL2; Tzourio-Mazoyer et al., 2002) using dorsal medial gyrus region (AAL2 number 5,6; Ma et al., 2018). Finally, we calculated the FC by correlating time-series of each network of interest (SAL, DLPFC), and the estimated connectivity value was inputted into the identical multiple regression model (see *equation 1*).

**Figure 1.**
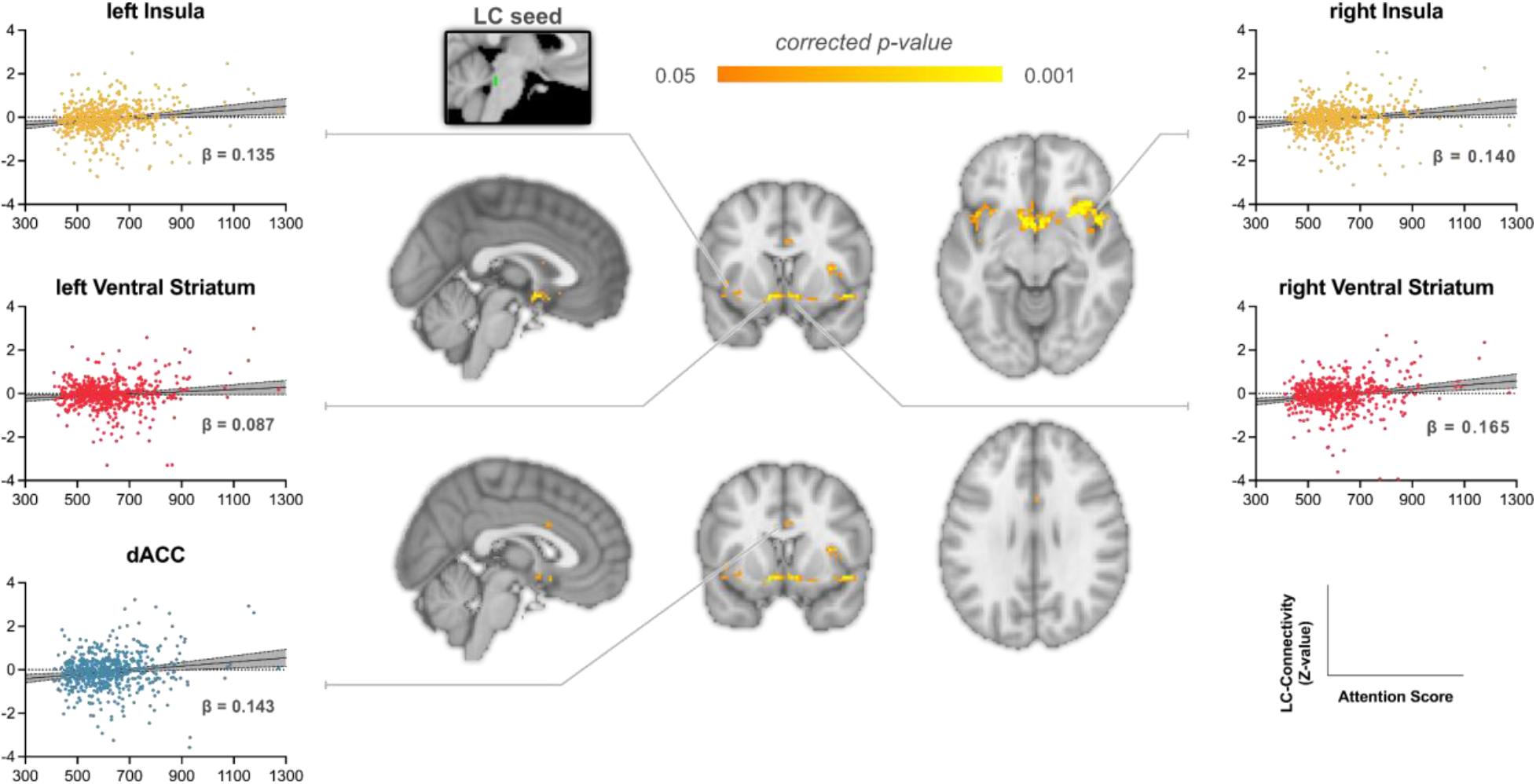
Significant regions resulting from multiple regression analyses illustrating regions associated with LC activity and the attention score. Scatter plots depict the relationship between the attention score and LC connectivity for illustration purposes. Beta represents standardized beta values.

### Effective Connectivity analysis between LC, Salience network and DLPFC

Finally, we examined the dynamics of functional connectivity between LC, SAL, and DLPFC, namely effective connectivity. To this end, we investigated effective connectivity patterns using granger causality (GC; Granger, 1969). Given that we did not assume an a priori model for the brain network relationships, this data-driven approach is an appropriate technique to investigate these relationships. We adopted pairwise-conditional GC within MVGC toolbox (Barnett & Seth, 2014) to diminish spurious effects of joint dependencies. Based on our result of previous section, the extracted time series from LC, SAL, and right DLPFC were used. The model order was selected as 2 using Bayesian information criteria. Individual’s estimated causality parameters were served as a dependent variable of the identical multiple regression model. Since there were six statistical analyses conducted, False Discovery Rate (FDR) correction was used (Benjamini & Hochberg, 1995).

## RESULTS

### LC Seed-Based Functional Connectivity

Multiple regression analyses of the LC seed-based connectivity revealed several significant regions within the salience network (Figure 1). Specifically, a significant association to the ANT score was found for connectivity between the LC and the bilateral insula, ventral striatum, and dorsal anterior cingulate cortex (dACC). Typically, the insula and dACC are regarded as core regions of the salience network; the ventral striatum is also included as an important subcortical area of the salience network (Seeley et al., 2007; Uddin, 2015). In our results, connectivity between these regions was positively associated with the ANT score, suggesting that increased LC-SAL communication may support attentional function (see Figure S1 Supplementary Materials).

We used the AAL2 atlas to cluster significant voxels into interpretable brain regions. In the AAL2, we adopted the defined insula regions intactly (region number 33, 34) but expanded certain boundaries for the dACC and the ventral striatum. For the dACC, we combined the dorsal part of ACC (35, 36) and most anterior parts of the mid cingulate cortex (37, 38). For the ventral striatum, caudate (75, 76) and dorsal parts of the olfactory area were combined (17, 18). Pearson correlations between the ANT scores and the averaged significant voxels within clusters revealed coefficients ranging between 0.09 to 0.17 (Table 1 and Figure 1). All significant voxels of LC connectivity associated with the ANT score are displayed in Figure S1 in Supplementary Materials.

**Table 1.**
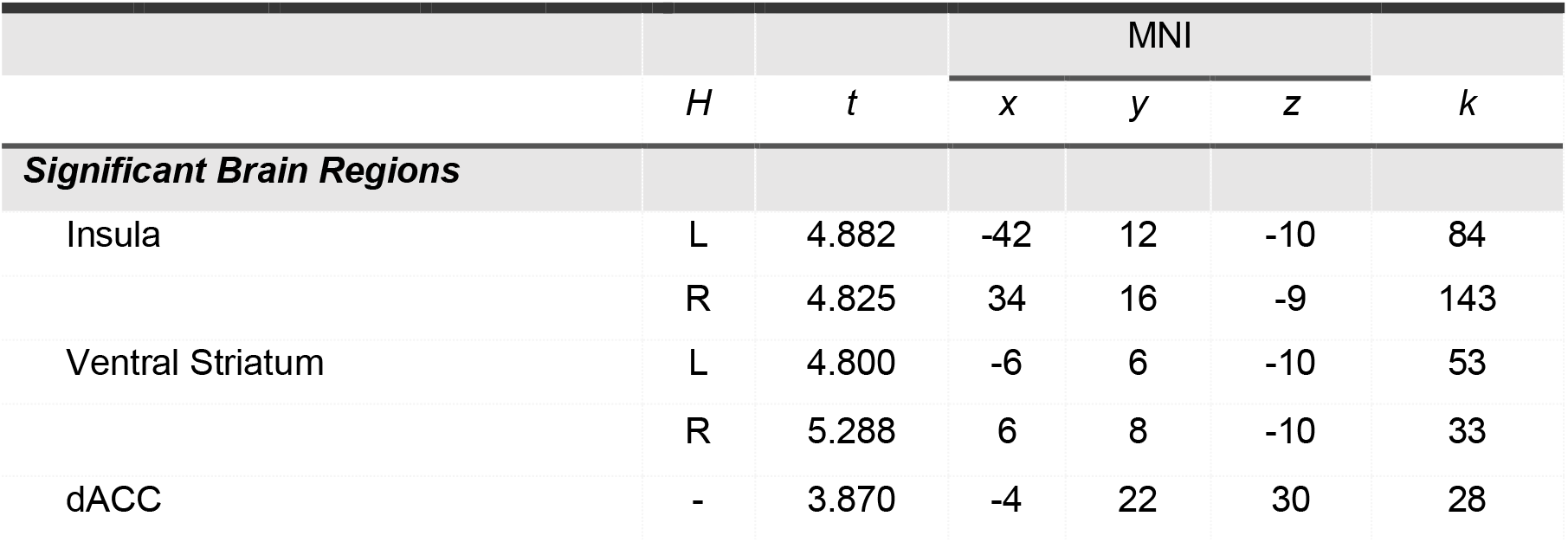
Significant brain regions associated with the ANT score on the LC seed-based whole-brain connectivity analysis. *H* = Hemisphere, *t* = *t*-value, *k* = number of significant voxels within the cluster, *r* = correlation coefficient with ANT score, *p* = p-value, Region labeling is based on the AAL2 atlas. The coordinate and *t*-value were based on the voxel whose maximum value.

**Table 2.**
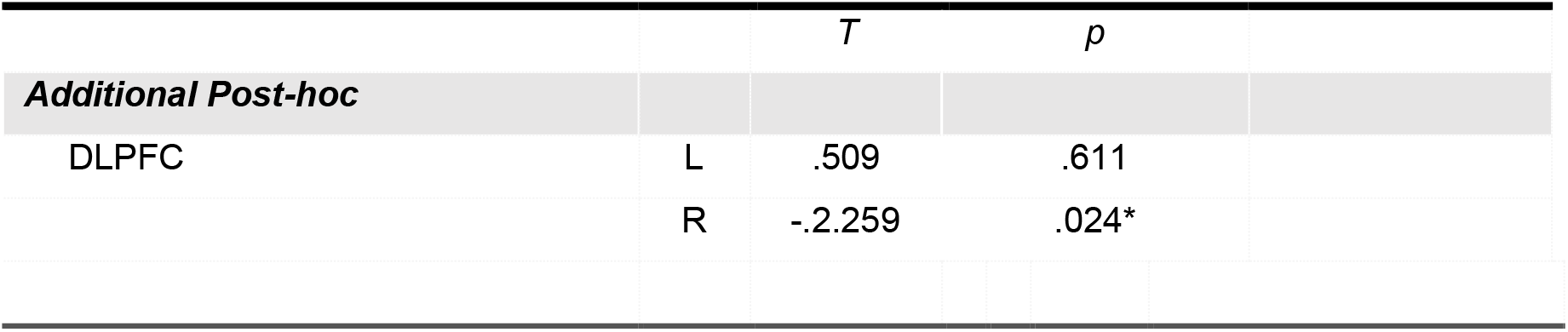
FC values for left and right hemispheres of DLPFC with SAL.

|  |  | *T* | *p* |
| --- | --- | --- | --- |
| **Additional Post-hoc** |  |  |  |
| DLPFC | L | .509 | .611 |
|  | R | -.2.259 | .024* |

### Salience network, attentional executive region and attention performance

Multiple regression analyses including SAL-DLPFC connectivity and the attention score, for each hemisphere, revealed that the connectivity between SAL-DLPFC is associated negatively with the ANT score in the right hemisphere (Table 1 and Figure 2). The association between the SAL-left DLPFC connectivity and the ANT score was not significant.

**Figure 2.**
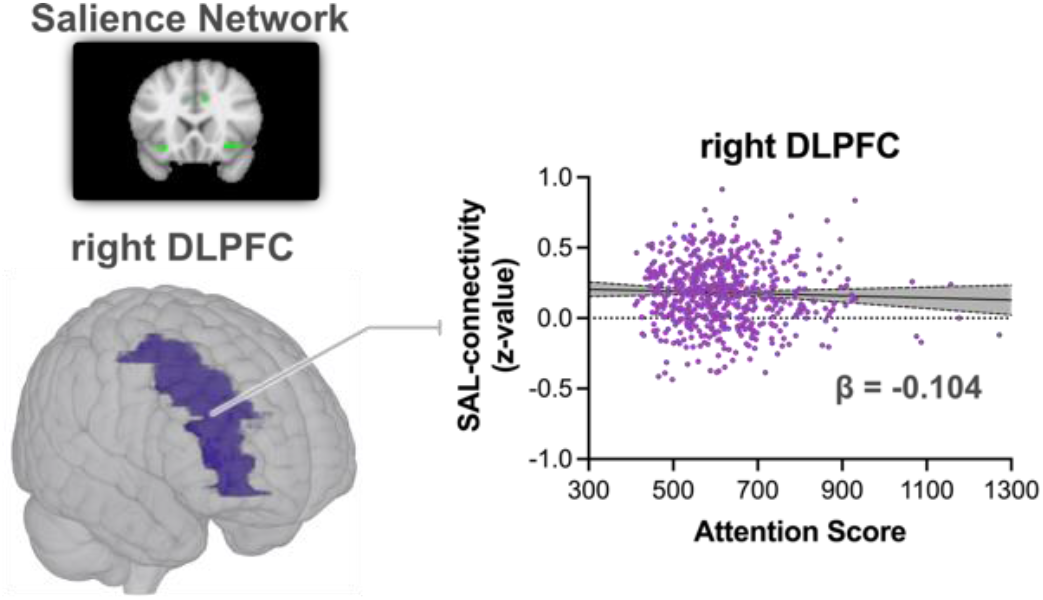
The association between the SAL – right DLPFC connectivity and the ANT score. Connectivity tended to decrease as the ANT score was increased. The DLPFC region was defined using dorsal medial gyrus in the AAL2 atlas. Beta represents the standardized beta values.

### Effective Connectivity analysis between LC, Salience network and DLPFC

The pairwise-conditional GC analysis showed two significant positive directional influences associative with the ANT score (Table 3 and Figure 3) The first flow was from SAL to LC (standardized *β* = .125, *t* = 2.695, *p* = .007, FDR-corrected *p* = .021), and the second was from right DLPFC to SAL (standardized *β* = .129, *t* = 2.804, *p* = .005, FDR-corrected *p* = .021).

**Table 3.** The pairwise-conditional granger causality parameters associated with the ANT score. *T* = *t*-value, *FDR corrected-p* = p-values corrected with false discovery rate (Benjamini & Hochberg, 1995).

| | | $T$ | $FDR\ corrected-p$ |
| --- | --- | --- | --- |
| *From* | *To* |  |  |
| SAL | right DLPFC | .123 | .974 |
| SAL | LC | 2.695 | .021* |
| right DLPFC | SAL | 2.804 | .021* |
| right DLPFC | LC | .033 | .974 |
| LC | SAL | .649 | .973 |
| LC | right DLPFC | 1.088 | .554 |

**Figure 3.**
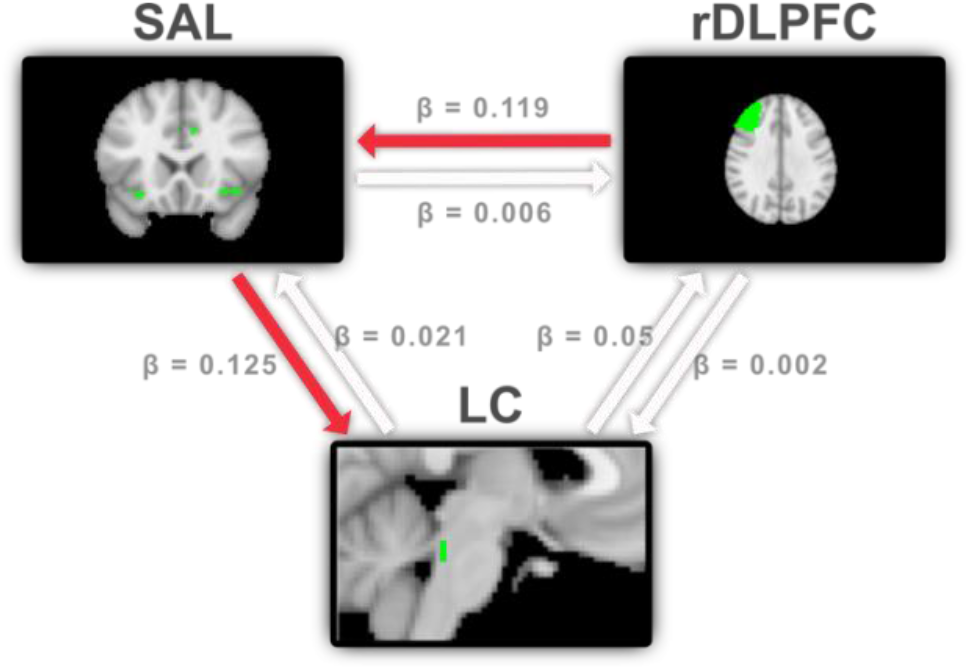
Effective connectivity results using a pairwise-conditional granger causality (GC) method. Two directed connectivity was associated with the ANT score significantly: from right DLPFC to SAL, and from SAL to the LC. Beta represents the standardized beta values. Red arrows indicate statistically significant at α = 0.05 with FDR correction.

## DISCUSSION

These findings support the application of the triple network model for neural cognitive function to the integration of other neural systems, in this case the LC specifically. Though often understood as acting as a neural switch for activation of the DMN and FPN, questions have remained regarding possible signaling inputs for the switching behavior itself, such as those from lower areas of the brain. With increases in connectivity between the LC and core regions of the SAL being associated with improved attentional performance, the possibility of an attentional signaling pathway beginning in the LC and traveling through the triple networks is evident. This expanded pathway has several advantages, including the LC’s prior associations with psychological arousal via norepinephrine release, as well as its links to neurological disorder and dysfunction. Critically, this integration of the LC into an attentional pathway represents a more specific signaling connectivity for the region to a connectivity network than has been previously identified.

Our additional analyses examining connectivity between the SAL and the DLPFC support a key theoretical finding relevant to another part of this model: the DLPFC represents a core region within the FPN network, and a negative relationship between LC-DLPFC connectivity and attention performance conforms to the concept of inattention through over-connection or over-activation of externalizing networks such as the FPN. This provides compelling support to the possibility that the extraneous processing of external stimuli, as represented by FPN activity in a setting with relatively low need (a resting state scan), represents a significant misapplication of neural and cognitive resources, which in turn lowers an individual’s overall attentive capabilities.

Additionally, the effective connectivity findings observed here suggest that the LC plays a pivotal role in sustained attention, and that the SAL may receive neural feedback or top-down control from other intrinsic network such as FPN (rDLPFC) and also signals to the LC to adjust arousal level. This is consistent with the view of that the SAL receives feedback from the cortex and its intrinsic networks, and sends input to the LC to alter arousal function within those networks (Unsworth & Robison, 2017). That is, the SAL changes the LC-NA system to increase the arousal level of the brain, and subsequently modulates the FPN activation level. However, the effective connectivity result should be interpreted with caution for several reasons. First, the current resting-state FC does not reflect task-related activity, and thus, we cannot rule out the possibility that the observed directionality between networks changes when participants are engaged in actual attentional processing. The bottom-up role of the LC may not have been prominent within in the current data. Second, the effective connectivity results do not necessarily indicate that the preponderance of neural communications begins from the FPN to the LC. The multiple regression only demonstrated that two pathways of directed connectivity were associated with attentional ability. Finally, the current GC analysis on the BOLD signal could not well capture the relatively slow effect of the LC-NA. For instance, Oyarzabal et al. (2022) showed that chemogenetic stimulation on tonic LC lead to NA release in other brain region with transient time in minutes (See Figure 4d in Oyarzabal et al., 2022). However, it is cautious as the current results are based on the resting-state connectivity when participants not actively engaged by the attentional processing. Instead, the current results suggest more about individual’s intrinsic foundation of attentional network that needs to be utilized during the actual attention performance. Future research might employ task-based assessments of LC activity related to the online attention process.

Nevertheless, these results suggest the possibility a top-down neural communication proportional to sustained attention. As described previously, a negative feedback signaling pathway would correspond with previous findings of network connections and with a regulated theory of optimal LC-NA signaling for attention performance. Given the specific absence of external stimuli due to the resting-state, such down-regulation could be expected. Additional support, particularly from alternative signaling patterns being observed when the FPN is in a task-positive state, is needed. Furthermore, the absence of a significant direct path in either direction between the DLPFC and LC continues to support the neural switching and processing role of the SAL, while also validating the pathway of the triple network model.

However, extant studies also imply that the SAL has been independently associated with abnormal functioning. For instance, SAL to FPN FC has been associated with attentive performance in task-based fMRI (Cai et al., 2021), and has also been identified as a differentiating neural pattern in attention deficit/hyperactivity disorder (ADHD) compared to non-pathological groups (Cai et al., 2019). Signaling from limbic regions to the SAL has also been identified as key differentiator of both ADHD diagnosis and symptom presentation (Damiani et al., 2021). SAL dysfunction has also been associated with inattention in other disorders, such as multiple sclerosis and ASD (Chen et al., 2021; Loitfelder et al., 2012). Age related differences in LC-SAL connectivity have also been observed in older adult samples (Lee et al., 2020; Liebe et al., 2022).

Attention, broadly defined as the process of devoting mental resources to given stimuli for specific periods of time, can be contextualized in a variety of individual processes and relationships. Arousal can be considered as both a requirement for and a component of attention, as it sets a broad capacity level for mental processes to operate within (Whyte, 1992). Attention is also related to the ability to perform sensory processing, such as visual or auditory processing (Woldorff et al., 1993). Attention is known for empowering executive control, which itself in turn resolves conflicts in attentional processes, allowing for more effective attentional processing and sustained attention (Posner & Fan, 2008). The focus of internal attention, meanwhile, has been observed to be related to the processing and recall of memory (Lewis-Peacock & Postle, 2012). Attention deficits extend across a number of disorders/conditions, such as obsessive-compulsive disorder, autism spectrum disorder, and attention-deficit/hyperactivity disorder (Biederman, 2005; Koch & Exner, 2015; New et al., 2010). The fact that attention may be divided into such discrete cognitive operations underlines that further research should work to better establish the associations between specific network connectivity operations and these respective cognitive processes.

Examining specific aspects of attention, as previously discussed, may require different study designs than those used here. The present analysis compared resting-state brain activity to the recorded performance on the ANT, a choice that compares the most generalizable neural data to a task that has been identified as likely to examine a given set of cognitive processes related to attention. Thus these findings depend somewhat upon the a priori assumptions of the task to develop resulting interpretation. One alternative design to examine cognitive aspects with fMRI would be the use of task-based fMRI. This would have the benefit of more directly instigating the cognitive processes of interest via targeted task design, and thereby observing neural data that is more directly comparable to actions intended within the task, such as attention. While useful for building theoretical frameworks between aspects of neural data (and activity) and cognitive processes, this specific attention-task fMRI does not facilitate a comparison to other outcome variables, such as behavioral reports or specific neuropsychological diagnoses. An approach with the most generalizable findings, then, would include intrinsic neural data, such as the resting-state, and relevant dependent clinical variables, with the relationship between the two supported by findings from previous study designs.

In principle, the identification of an attentional processing pathway suggests additional associations with other dysfunctions related to attention deficits. Identifying a consistent neural pathway associated with attention deficits would offer profound implications across diagnostic concepts and disorder categories. Attention deficits and LC dysfunction have been identified in several neurodegenerative disorders including ADHD, OCD, and PTSD. The current study may provide an alternative model of psychopathology, one based on the transdiagnostic dispersion of dysfunctions and relationships between the LC, SAL and FPN, and could ultimately help to establish a general neural model for attention deficits. The Hierarchical Taxonomy of Psychopathology (HiTOP; Kotov et al., 2017), one of the leading approaches to this methodology, has not at present established neural or cognitive constructs for neurodevelopmental disorders. Rdoc (Insel et al., 2010) is a research-focused matrix encompassing this principle of transdiagnostic symptom identification. Identifying connections between neural pathways such as a LC-triple network model and cognitive constructs such as attention dysfunction will prove vital in translating across the empirical observation of symptoms and more traditional disorder nosologies.

Moving forward, studies may use multiple methods for extending and expanding the findings here given the present study contain a number of limitations, mainly derived from using previously collected public dataset. First, the small size of the LC as a region generates substantial challenges in deriving signal purely from it, and though several techniques have been established to improve the signal extraction, the preparatory work required for these meant they were not available for the present analysis. Spatial resolution of the LC beyond the standard 3T structural image (e.g., 7T) is one method to improve the extraction process (Morris et al., 2020). Additionally, specializing additional T1 images with a decreased field of view to increase spatial resolution within the brainstem itself is another option (e.g., Clewett et al., 2016, 2018). Regarding the BOLD signal of the LC, it should be improved by the collection of physiological data, especially cardiovascular information such as heart rate and blood pulse, allowing us to separate LC signal from physio noise. Importantly, recent studies showed that ICA-denoising approach adopted in the current study (i.e., ICA-AROMA) reliably separate physiological and motion noise and improve LC BOLD signal (e.g., Lee et al., 2018, 2020; Prokopiou et al., 2022).

Future studies should also seek to examine more specific populations. The LC has been noted to exhibit different activation patterns across the lifespan (Song et al., 2021).As described earlier, developmental theories of neural pathologies are associated with differential expectations regarding neural activity at different ages. Analyses of more granularly defined age populations may, in turn, reveal a great deal in terms of how these connections and associated performance change across the lifespan. In particular, studies containing child and adolescent populations may be able to better discern developmental processes within the LC and accompanying connectivity networks. In combination with clinical populations, this could greatly inform the underlying neural mechanisms of the pathology of attention deficits as a specific symptom within syndromes/disorders, with important implications for the identification and treatment of these disorders more generally.

Previous findings within neuroimaging studies of attention have suggested that dysfunctional network switching between the SAL and the network activating and regulating processes of the LC may present a unified pathway. The positive relationship identified here between LC-SAL functional connectivity and attentional performance represents an exciting initial step towards establishing direct evidence of such a relationship. The post-hoc evaluations of network connections to the SAL also support the principles of the triple network model as applied to a performative attention outcome. Subsequent research should attempt to more directly and fully demonstrate this relationship.

## Acknowledgements

We would like to thank the Neuroimaging Informatics Tools and Resources Clearinghouse and all of the researchers who have contributed with data to the 1000 Functional Connectomes Project and to the Enhanced Nathan Kline Institute - Rockland Sample.

## Data Availability

Source Data: https://osf.io/5bwr8/

## References

Aertsen, A. M., Gerstein, G. L., Habib, M. K., & Palm, G. (1989). Dynamics of neuronal firing correlation: Modulation of “effective connectivity.” Journal of Neurophysiology, 61(5), 900–917. https://doi.org/10.1152/jn.1989.61.5.900

Alakörkkö, T., Saarimäki, H., Glerean, E., Saramäki, J., & Korhonen, O. (2017). Effects of spatial smoothing on functional brain networks. European Journal of Neuroscience, 46(9), 2471–2480. https://doi.org/10.1111/ejn.13717

Aston-Jones, G., & Cohen, J. D. (2005). An integrative theory of locus coeruleus-norepinephrine function: Adaptive gain and optimal performance. Annu. Rev. Neurosci., 28, 403–450.

Avants, B. B., Tustison, N. J., Stauffer, M., Song, G., Wu, B., & Gee, J. C. (2014). The Insight ToolKit image registration framework. Frontiers in Neuroinformatics, 8. https://doi.org/10.3389/fninf.2014.00044

Barnett, L., & Seth, A. K. (2014). The MVGC multivariate Granger causality toolbox: A new approach to Granger-causal inference. Journal of Neuroscience Methods, 223, 50–68. https://doi.org/10.1016/j.jneumeth.2013.10.018

Benjamini, Y., & Hochberg, Y. (1995). Controlling the False Discovery Rate: A Practical and Powerful Approach to Multiple Testing. Journal of the Royal Statistical Society: Series B (Methodological), 57(1), 289–300. https://doi.org/10.1111/j.2517-6161.1995.tb02031.x

Biederman, J. (2005). Attention-deficit/hyperactivity disorder: A selective overview. Biological Psychiatry, 57(11), 1215–1220. https://doi.org/10.1016/j.biopsych.2004.10.020

Biswal, B., Zerrin Yetkin, F., Haughton, V. M., & Hyde, J. S. (1995). Functional connectivity in the motor cortex of resting human brain using echo-planar mri. Magnetic Resonance in Medicine, 34(4), 537–541. https://doi.org/10.1002/mrm.1910340409

Bonnelle, V., Ham, T. E., Leech, R., Kinnunen, K. M., Mehta, M. A., Greenwood, R. J., & Sharp, D. J. (2012). Salience network integrity predicts default mode network function after traumatic brain injury. Proceedings of the National Academy of Sciences, 109(12), 4690–4695. https://doi.org/10.1073/pnas.1113455109

Buckner, R. L., & Vincent, J. L. (2007). Unrest at rest: Default activity and spontaneous network correlations. NeuroImage, 37(4), 1091–1096. https://doi.org/10.1016/j.neuroimage.2007.01.010

Cai, W., Griffiths, K., Korgaonkar, M. S., Williams, L. M., & Menon, V. (2021). Inhibition-related modulation of salience and frontoparietal networks predicts cognitive control ability and inattention symptoms in children with ADHD. Molecular Psychiatry, 26(8), 4016–4025. https://doi.org/10.1038/s41380-019-0564-4

Cai, W., Griffiths, K., Korgaonkar, M., Williams, L., & Menon, V. (2019). F56. Task-Evoked Effective Connectivity in Salience and Central Executive Networks Predicts Cognitive Control Ability and Inattention Symptoms in Children With ADHD. Biological Psychiatry, 85(10), S234–S235. https://doi.org/10.1016/j.biopsych.2019.03.593

Chand, G. B., & Dhamala, M. (2017). Interactions between the anterior cingulate-insula network and the fronto-parietal network during perceptual decision-making. NeuroImage, 152, 381–389. https://doi.org/10.1016/j.neuroimage.2017.03.014

Chen, Y.-Y., Uljarevic, M., Neal, J., Greening, S., Yim, H., & Lee, T.-H. (2021). Excessive functional coupling with less variability between salience and default-mode networks in Autism Spectrum Disorder. Biological Psychiatry: Cognitive Neuroscience and Neuroimaging. https://doi.org/10.1016/j.bpsc.2021.11.016

Clewett, D. V., Huang, R., Velasco, R., Lee, T. H., & Mather, M. (2018). Locus coeruleus activity strengthens prioritized memories under arousal. Journal of Neuroscience, 38(6), 1558–1574. https://doi.org/10.1523/JNEUROSCI.2097-17.2017

Clewett, D. V., Lee, T.-H., Greening, S., Ponzio, A., Margalit, E., & Mather, M. (2016). Neuromelanin marks the spot: Identifying a locus coeruleus biomarker of cognitive reserve in healthy aging. Neurobiology of Aging, 37, 117–126. https://doi.org/10.1016/j.neurobiolaging.2015.09.019

Dahlöf, C., Engberg, G., & Svensson, T. H. (1981). Effects of β-adrenoceptor antagonists on the firing rate of noradrenergic neurones in the locus coeruleus of the rat. Naunyn-Schmiedeberg’s Archives of Pharmacology, 317(1), 26–30.

Damiani, S., Tarchi, L., Scalabrini, A., Marini, S., Provenzani, U., Rocchetti, M., Oliva, F., & Politi, P. (2021). Beneath the surface: Hyper-connectivity between caudate and salience regions in ADHD fMRI at rest. European Child & Adolescent Psychiatry, 30(4), 619–631. https://doi.org/10.1007/s00787-020-01545-0

Devilbiss, D. M., & Waterhouse, B. D. (2011). Phasic and tonic patterns of locus coeruleus output differentially modulate sensory network function in the awake rat. Journal of Neurophysiology, 105(1), 69–87. https://doi.org/10.1152/jn.00445.2010

Devoto, P., Flore, G., Saba, P., Fa, M., & Gessa, G. L. (2005). Stimulation of the locus coeruleus elicits noradrenaline and dopamine release in the medial prefrontal and parietal cortex. Journal of Neurochemistry, 92(2), 368–374. https://doi.org/10.1111/j.1471-4159.2004.02866.x

Fan, J., McCandliss, B. D., Sommer, T., Raz, A., & Posner, M. I. (2002). Testing the Efficiency and Independence of Attentional Networks. Journal of Cognitive Neuroscience, 14(3), 340–347. https://doi.org/10.1162/089892902317361886

Fortenbaugh, F. C., DeGutis, J., & Esterman, M. (2017). Recent theoretical, neural, and clinical advances in sustained attention research: Sustained attention. Annals of the New York Academy of Sciences, 1396(1), 70–91. https://doi.org/10.1111/nyas.13318

Friston, K. J., Frith, C. D., Liddle, P. F., & Frackowiak, R. S. (1993). Functional connectivity: The principal-component analysis of large (PET) data sets. Journal of Cerebral Blood Flow & Metabolism, 13(1), 5–14.

Galvao-Carmona, A., GonzÃilez-Rosa, J. J., Hidalgo-MuÃ±oz, A. R., PÃiramo, D., BenÃ-tez, M. L., Izquierdo, G., & VÃzquez-Marrufo, M. (2014). Disentangling the attention network test: Behavioral, event related potentials, and neural source analyses. Frontiers in Human Neuroscience, 8. https://doi.org/10.3389/fnhum.2014.00813

Goulden, N., Khusnulina, A., Davis, N. J., Bracewell, R. M., Bokde, A. L., McNulty, J. P., & Mullins, P. G. (2014). The salience network is responsible for switching between the default mode network and the central executive network: Replication from DCM. NeuroImage, 99, 180–190. https://doi.org/10.1016/j.neuroimage.2014.05.052

Granger, C. W. (1969). Investigating causal relations by econometric models and cross-spectral methods. Econometrica: Journal of the Econometric Society, 424–438.

Gürsel, D. A., Avram, M., Sorg, C., Brandl, F., & Koch, K. (2018). Frontoparietal areas link impairments of large-scale intrinsic brain networks with aberrant fronto-striatal interactions in OCD: A meta-analysis of resting-state functional connectivity. Neuroscience & Biobehavioral Reviews, 87, 151–160. https://doi.org/10.1016/j.neubiorev.2018.01.016

Huang, Y., Yu, S., Wilson, G., Park, J., Cheng, M., Kong, X., Lu, T., & Kong, J. (2021). Altered Extended Locus Coeruleus and Ventral Tegmental Area Networks in Boys with Autism Spectrum Disorders: A Resting-State Functional Connectivity Study. Neuropsychiatric Disease and Treatment, 10.

Insel, T., Cuthbert, B., Garvey, M., Heinssen, R., Pine, D. S., Quinn, K., Sanislow, C., & Wang, P. (2010). Research Domain Criteria (RDoC): Toward a New Classification Framework for Research on Mental Disorders. American Journal of Psychiatry, 167(7), 748–751. https://doi.org/10.1176/appi.ajp.2010.09091379

Jacobs, H. I. L., Wiese, S., van de Ven, V., Gronenschild, E. H. B. M., Verhey, F. R. J., & Matthews, P. M. (2015). Relevance of parahippocampal-locus coeruleus connectivity to memory in early dementia. Neurobiology of Aging, 36(2), 618–626. https://doi.org/10.1016/j.neurobiolaging.2014.10.041

Jodoj, E., Chiang, C., & Aston-Jones, G. (1998). Potent excitatory influence of prefrontal cortex activity on noradrenergic locus coeruleus neurons. Neuroscience, 83(1), 63–79. https://doi.org/10.1016/S0306-4522(97)00372-2

Johnson, K. A., Robertson, I. H., Barry, E., Mulligan, A., Dáibhis, A., Daly, M., Watchorn, A., Gill, M., & Bellgrove, M. A. (2008). Impaired conflict resolution and alerting in children with ADHD: Evidence from the Attention Network Task (ANT). Journal of Child Psychology and Psychiatry, 49(12), 1339–1347. https://doi.org/10.1111/j.1469-7610.2008.01936.x

Jung, J., Lambon Ralph, M. A., & Jackson, R. L. (2022). Subregions of DLPFC Display Graded yet Distinct Structural and Functional Connectivity. The Journal of Neuroscience, 42(15), 3241–3252. https://doi.org/10.1523/JNEUROSCI.1216-21.2022

Keller, J. B., Hedden, T., Thompson, T. W., Anteraper, S. A., Gabrieli, J. D. E., & Whitfield-Gabrieli, S. (2015). Resting-state anticorrelations between medial and lateral prefrontal cortex: Association with working memory, aging, and individual differences. Cortex, 64, 271–280. https://doi.org/10.1016/j.cortex.2014.12.001

Keren, N. I., Lozar, C. T., Harris, K. C., Morgan, P. S., & Eckert, M. A. (2009). In vivo mapping of the human locus coeruleus. NeuroImage, 47(4), 1261–1267. https://doi.org/10.1016/j.neuroimage.2009.06.012

Koch, J., & Exner, C. (2015). Selective attention deficits in obsessive–compulsive disorder: The role of metacognitive processes. Psychiatry Research, 225(3), 550–555. https://doi.org/10.1016/j.psychres.2014.11.049

Kotov, R., Krueger, R. F., Watson, D., Achenbach, T. M., Althoff, R. R., Bagby, R. M., Brown, T. A., Carpenter, W. T., Caspi, A., Clark, L. A., Eaton, N. R., Forbes, M. K., Forbush, K. T., Goldberg, D., Hasin, D., Hyman, S. E., Ivanova, M. Y., Lynam, D. R., Markon, K., … Zimmerman, M. (2017). The Hierarchical Taxonomy of Psychopathology (HiTOP): A dimensional alternative to traditional nosologies. Journal of Abnormal Psychology, 126(4), 454–477. https://doi.org/10.1037/abn0000258

Laird, A. R., Fox, P. M., Eickhoff, S. B., Turner, J. A., Ray, K. L., McKay, D. R., Glahn, D. C., Beckmann, C. F., Smith, S. M., & Fox, P. T. (2011). Behavioral Interpretations of Intrinsic Connectivity Networks. Journal of Cognitive Neuroscience, 23(12), 4022–4037. https://doi.org/10.1162/jocn_a_00077

Landis, D., Courtney, W., Dieringer, C., Kelly, R., King, M., Miller, B., Wang, R., Wood, D., Turner, J. A., & Calhoun, V. D. (2016). COINS Data Exchange: An open platform for compiling, curating, and disseminating neuroimaging data. NeuroImage, 124, 1084–1088. https://doi.org/10.1016/j.neuroimage.2015.05.049

Lee, T.-H., Greening, S. G., Ueno, T., Clewett, D., Ponzio, A., Sakaki, M., & Mather, M. (2018). Arousal increases neural gain via the locus coeruleus–noradrenaline system in younger adults but not in older adults. Nature Human Behaviour, 2(5), 356–366. https://doi.org/10.1038/s41562-018-0344-1

Lee, T.-H., Kim, S. H., Katz, B., & Mather, M. (2020). The Decline in Intrinsic Connectivity Between the Salience Network and Locus Coeruleus in Older Adults: Implications for Distractibility. Frontiers in Aging Neuroscience, 12, 2–2.

Lewis-Peacock, J. A., & Postle, B. R. (2012). Decoding the internal focus of attention. Neuropsychologia, 50(4), 470–478. https://doi.org/10.1016/j.neuropsychologia.2011.11.006

Liebe, T., Dordevic, M., Kaufmann, J., Avetisyan, A., Skalej, M., & Müller, N. (2022). Investigation of the functional pathogenesis of mild cognitive impairment by localisation-based locus coeruleus resting-state FMRI. Human Brain Mapping, hbm.26039. https://doi.org/10.1002/hbm.26039

Loitfelder, M., Filippi, M., Rocca, M., Valsasina, P., Ropele, S., Jehna, M., Fuchs, S., Schmidt, R., Neuper, C., Fazekas, F., & Enzinger, C. (2012). Abnormalities of Resting State Functional Connectivity Are Related to Sustained Attention Deficits in MS. PLoS ONE, 7(8), e42862. https://doi.org/10.1371/journal.pone.0042862

Lowe, M. J., Dzemidzic, M., Lurito, J. T., Mathews, V. P., & Phillips, M. D. (2000). Correlations in low-frequency BOLD fluctuations reflect cortico-cortical connections. NeuroImage, 12(5), 582–587. https://doi.org/10.1006/nimg.2000.0654

Luna, F. G., Román-Caballero, R., Barttfeld, P., Lupiáñez, J., & Martín-Arévalo, E. (2020). A High-Definition tDCS and EEG study on attention and vigilance: Brain stimulation mitigates the executive but not the arousal vigilance decrement. Neuropsychologia, 142, 107447. https://doi.org/10.1016/j.neuropsychologia.2020.107447

Ma, L., Steinberg, J. L., Bjork, J. M., Keyser-Marcus, L., Vassileva, J., Zhu, M., Ganapathy, V., Wang, Q., Boone, E. L., Ferré, S., Bickel, W. K., & Gerard Moeller, F. (2018). Fronto-striatal effective connectivity of working memory in adults with cannabis use disorder. Psychiatry Research: Neuroimaging, 278, 21–34. https://doi.org/10.1016/j.pscychresns.2018.05.010

MacLeod, J. W., Lawrence, M. A., McConnell, M. M., Eskes, G. A., Klein, R. M., & Shore, D. I. (2010). Appraising the ANT: Psychometric and theoretical considerations of the Attention Network Test. Neuropsychology, 24(5), 637–651. https://doi.org/10.1037/a0019803

Marzo, A., Totah, N. K., Neves, R. M., Logothetis, N. K., & Eschenko, O. (2014). Unilateral electrical stimulation of rat locus coeruleus elicits bilateral response of norepinephrine neurons and sustained activation of medial prefrontal cortex. Journal of Neurophysiology, 111(12), 2570–2588. https://doi.org/10.1152/jn.00920.2013

Matchett, B. J., Grinberg, L. T., Theofilas, P., & Murray, M. E. (2021). The mechanistic link between selective vulnerability of the locus coeruleus and neurodegeneration in Alzheimer’s disease. Acta Neuropathologica, 141, 20.

McConnell, M. M., & Shore, D. I. (2011). Mixing measures: Testing an assumption of the attention network test. Attention, Perception, & Psychophysics, 73(4), 1096–1107. https://doi.org/10.3758/s13414-010-0085-3

Menon, V. (2011). Large-scale brain networks and psychopathology: A unifying triple network model. Trends in Cognitive Sciences, 15(10), 483–506. https://doi.org/10.1016/j.tics.2011.08.003

Morris, L. S., Tan, A., Smith, D. A., Grehl, M., Han-Huang, K., Naidich, T. P., Charney, D. S., Balchandani, P., Kundu, P., & Murrough, J. W. (2020). Sub-millimeter variation in human locus coeruleus is associated with dimensional measures of psychopathology: An in vivo ultra-high field 7-Tesla MRI study. NeuroImage: Clinical, 25, 102148. https://doi.org/10.1016/j.nicl.2019.102148

Naegeli, C., Zeffiro, T., Piccirelli, M., Jaillard, A., Weilenmann, A., Hassanpour, K., Schick, M., Rufer, M., Mueller-Orr, S., & Pfeiffer, C. (2018). Locus Coeruleus Activity Mediates Hyperresponsiveness in Posttraumatic Stress Disorder. Biological Psychiatry, 9.

Nai-shin, C., & E, B. F. (1973). Norepinephrine-Containing Neurons: Changes in Spontaneous Discharge Patterns during Sleeping and Waking. Science, 179(4076), 908–910. https://doi.org/10.1126/science.179.4076.908

Nejati, V., Salehinejad, M. A., & Nitsche, M. A. (2018). Interaction of the Left Dorsolateral Prefrontal Cortex (l-DLPFC) and Right Orbitofrontal Cortex (OFC) in Hot and Cold Executive Functions: Evidence from Transcranial Direct Current Stimulation (tDCS). Neuroscience, 369, 109–123. https://doi.org/10.1016/j.neuroscience.2017.10.042

New, J. J., Schultz, R. T., Wolf, J., Niehaus, J. L., Klin, A., German, T. C., & Scholl, B. J. (2010). The scope of social attention deficits in autism: Prioritized orienting to people and animals in static natural scenes. Neuropsychologia, 48(1), 51–59. https://doi.org/10.1016/j.neuropsychologia.2009.08.008

Niendam, T. A., Laird, A. R., Ray, K. L., Dean, Y. M., Glahn, D. C., & Carter, C. S. (2012). Meta-analytic evidence for a superordinate cognitive control network subserving diverse executive functions. Cognitive, Affective, & Behavioral Neuroscience, 12(2), 241–268. https://doi.org/10.3758/s13415-011-0083-5

Nooner, K. B., Colcombe, S. J., Tobe, R. H., Mennes, M., Benedict, M. M., Moreno, A. L., Panek, L. J., Brown, S., Zavitz, S. T., Li, Q., Sikka, S., Gutman, D., Bangaru, S., Schlachter, R. T., Kamiel, S. M., Anwar, A. R., Hinz, C. M., Kaplan, M. S., Rachlin, A. B., … Milham, M. P. (2012). The NKI-Rockland Sample: A Model for Accelerating the Pace of Discovery Science in Psychiatry. Frontiers in Neuroscience, 6. https://doi.org/10.3389/fnins.2012.00152

Oyarzabal, E. A., Hsu, L.-M., Das, M., Chao, T.-H. H., Zhou, J., Song, S., Zhang, W., Smith, K. G., Sciolino, N. R., Evsyukova, I. Y., Yuan, H., Lee, S.-H., Cui, G., Jensen, P., & Shih, Y.-Y. I. (2022). Chemogenetic stimulation of tonic locus coeruleus activity strengthens the default mode network. Science Advances, 8(17), eabm9898. https://doi.org/10.1126/sciadv.abm9898

Posner, M. I., & Fan, J. (2008). Attention as an organ system. Topics in Integrative Neuroscience, 31.

Prokopiou, P. C., Engels-Domínguez, N., Papp, K. V., Scott, M. R., Schultz, A. P., Schneider, C., Farrell, M. E., Buckley, R. F., Quiroz, Y. T., El Fakhri, G., Rentz, D. M., Sperling, R. A., Johnson, K. A., & Jacobs, H. I. L. (2022). Lower novelty-related locus coeruleus function is associated with Aβ-related cognitive decline in clinically healthy individuals. Nature Communications, 13(1), 1571. https://doi.org/10.1038/s41467-022-28986-2

Pruim, R. H. R., Mennes, M., van Rooij, D., Llera, A., Buitelaar, J. K., & Beckmann, C. F. (2015). ICA-AROMA: A robust ICA-based strategy for removing motion artifacts from fMRI data. NeuroImage, 112, 267–277. https://doi.org/10.1016/j.neuroimage.2015.02.064

Rajkowski, J. (2000). Prominent projections from the anterior cingulate cortex to the locus coeruleus (LC) in rhesus monkey. 26, 2230.

Sara, S. J. (2009). The locus coeruleus and noradrenergic modulation of cognition. Nature Reviews Neuroscience, 10(3), 211–223. https://doi.org/10.1038/nrn2573

Sarter, M., Givens, B., & Bruno, J. P. (2001). The cognitive neuroscience of sustained attention: Where top-down meets bottom-up (Brain Research Reviews, Vol. 35, pp. 146–160). https://www.elsevier.com/locate/bres

Seeley, W. W., Menon, V., Schatzberg, A. F., Keller, J., Glover, G. H., Kenna, H., Reiss, A. L., & Greicius, M. D. (2007). Dissociable Intrinsic Connectivity Networks for Salience Processing and Executive Control. Journal of Neuroscience, 27(9), 2349–2356. https://doi.org/10.1523/JNEUROSCI.5587-06.2007

Smith, S., & Nichols, T. (2009). Threshold-free cluster enhancement: Addressing problems of smoothing, threshold dependence and localisation in cluster inference. NeuroImage, 44(1), 83–98. https://doi.org/10.1016/j.neuroimage.2008.03.061

Song, I., Neal, J., & Lee, T.-H. (2021). Age-Related Intrinsic Functional Connectivity Changes of Locus Coeruleus from Childhood to Older Adults. Brain Sciences, 11, 1485–1485. https://doi.org/10.3390/brainsci11111485

Thiele, A., & Bellgrove, M. A. (2018). Neuromodulation of Attention. Neuron, 97(4), 769–785. https://doi.org/10.1016/j.neuron.2018.01.008

Tobe, R. H., MacKay-Brandt, A., Lim, R., Kramer, M., Breland, M. M., Tu, L., Tian, Y., Trautman, K. D., Hu, C., Sangoi, R., Alexander, L., Gabbay, V., Castellanos, F. X., Leventhal, B. L., Craddock, R. C., Colcombe, S. J., Franco, A. R., & Milham, M. P. (2022). A longitudinal resource for studying connectome development and its psychiatric associations during childhood. Scientific Data, 9(1), 300. https://doi.org/10.1038/s41597-022-01329-y

Tzourio-Mazoyer, N., Landeau, B., Papathanassiou, D., Crivello, F., Etard, O., Delcroix, N., Mazoyer, B., & Joliot, M. (2002). Automated Anatomical Labeling of Activations in SPM Using a Macroscopic Anatomical Parcellation of the MNI MRI Single-Subject Brain. NeuroImage, 15(1), 273–289. https://doi.org/10.1006/nimg.2001.0978

Uddin, L. Q. (2015). Salience processing and insular cortical function and dysfunction. Nature Reviews Neuroscience, 16(1), 55–61. https://doi.org/10.1038/nrn3857

Unsworth, N., & Robison, M. K. (2017). A locus coeruleus-norepinephrine account of individual differences in working memory capacity and attention control. Psychonomic Bulletin & Review, 24(4), 1282–1311. https://doi.org/10.3758/s13423-016-1220-5

van den Hurk, P. A. M., Giommi, F., Gielen, S. C., Speckens, A. E. M., & Barendregt, H. P. (2010). Greater efficiency in attentional processing related to mindfulness meditation. Quarterly Journal of Experimental Psychology, 63(6), 1168–1180. https://doi.org/10.1080/17470210903249365

Vazey, E. M., Moorman, D. E., & Aston-Jones, G. (2018). Phasic locus coeruleus activity regulates cortical encoding of salience information. Proceedings of the National Academy of Sciences of the United States of America, 115(40), E9439–E9448. https://doi.org/10.1073/pnas.1803716115

Wang, J., Wang, Y., Huang, H., Jia, Y., Zheng, S., Zhong, S., Chen, G., Huang, L., & Huang, R. (2020). Abnormal dynamic functional network connectivity in unmedicated bipolar and major depressive disorders based on the triple-network model. Psychological Medicine, 50(3), 465–474. https://doi.org/10.1017/S003329171900028X

Wei, H., An, J., Shen, H., Zeng, L. L., Qiu, S., & Hu, D. (2016). Altered effective connectivity among core neurocognitive networks in idiopathic generalized epilepsy: An fMRI evidence. Frontiers in Human Neuroscience, 10(SEP2016). https://doi.org/10.3389/fnhum.2016.00447

Whyte, J. (1992). Attention and arousal: Basic science aspects. Archives of Physical Medicine and Rehabilitation, 73(10), 940–949. https://doi.org/10.5555/uri:pii:000399939290266Y

Winkler, A. M., Ridgway, G. R., Webster, M. A., Smith, S. M., & Nichols, T. E. (2014). Permutation inference for the general linear model. NeuroImage, 92, 381–397. https://doi.org/10.1016/j.neuroimage.2014.01.060

Woldorff, M. G., Gallen, C. C., Hampson, S. A., Hillyard, S. A., Pantev, C., Sobel, D., & Bloom, F. E. (1993). Modulation of early sensory processing in human auditory cortex during auditory selective attention. Proceedings of the National Academy of Sciences, 90(18), 8722–8722. https://doi.org/10.1073/pnas.90.18.8722

Yu, E., Liao, Z., Tan, Y., Qiu, Y., Zhu, J., Han, Z., Wang, J., Wang, X., Wang, H., Chen, Y., Zhang, Q., Li, Y., Mao, D., & Ding, Z. (2019). High-sensitivity neuroimaging biomarkers for the identification of amnestic mild cognitive impairment based on resting-state fMRI and a triple network model. Brain Imaging and Behavior, 13(1), 1–14. https://doi.org/10.1007/s11682-017-9727-6

Zerbi, V., Floriou-Servou, A., Markicevic, M., Vermeiren, Y., Sturman, O., Privitera, M., von Ziegler, L., Ferrari, K. D., Weber, B., De Deyn, P. P., Wenderoth, N., & Bohacek, J. (2019). Rapid Reconfiguration of the Functional Connectome after Chemogenetic Locus Coeruleus Activation. Neuron, 103(4), 702–718.e5. https://doi.org/10.1016/j.neuron.2019.05.034

