## Supplemental Information for "Association of intrinsic functional connectivity between the locus coeruleus and salience network with attentional ability"

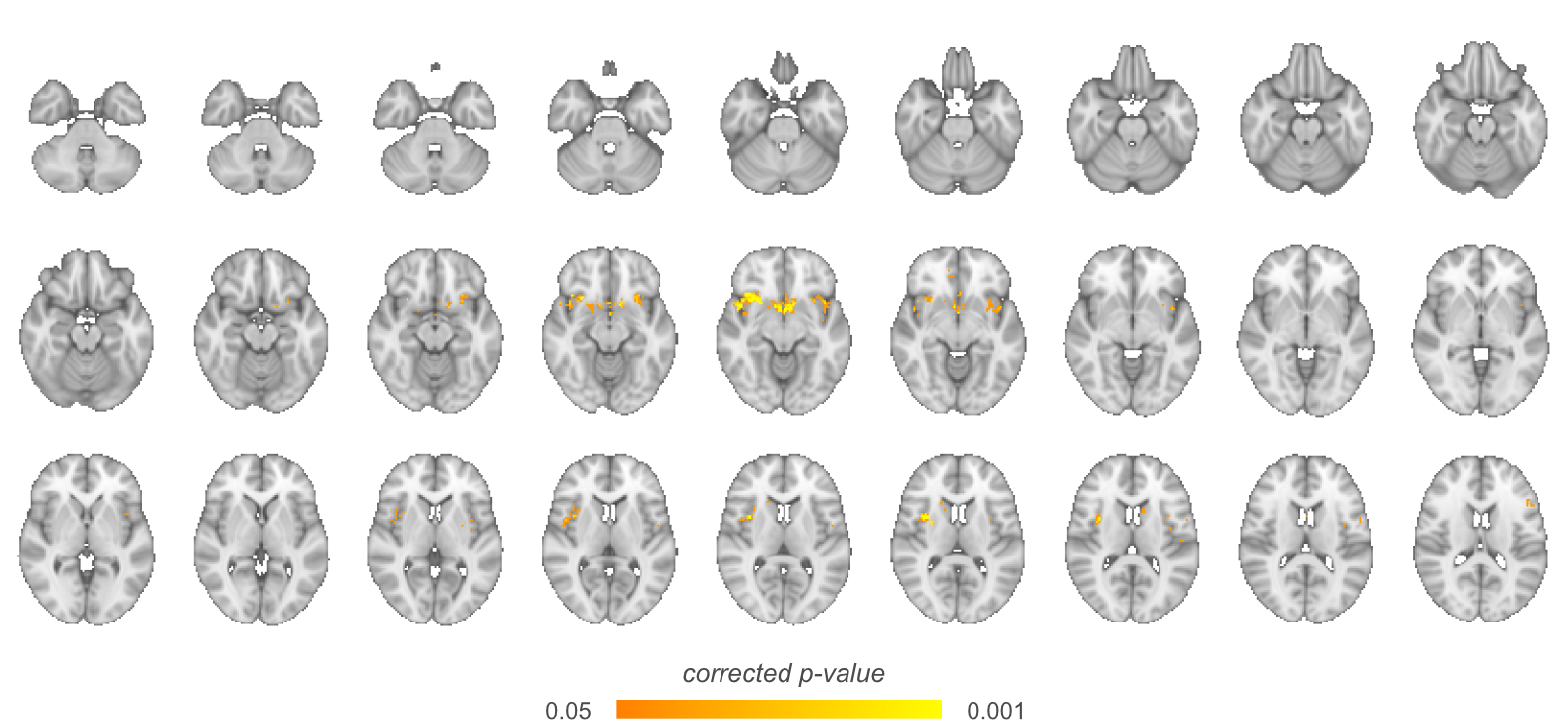

*Supplementary Figure 1.* The complete collection of regions with LC FC positively associated with the attention score.
